# Dynamic interplay of innate and adaptive immunity during sterile retinal inflammation: Insights from the transcriptome

**DOI:** 10.1101/267757

**Authors:** Riccardo Natoli, Elizabeth Mason, Haihan Jiao, Aaron Chuah, Hardip Patel, Nilisha Fernando, Krisztina Valter, Christine A. Wells, Jan Provis, Matt Rutar

## Abstract

The pathogenesis of many retinal degenerations, such as age-related macular degeneration (AMD), is punctuated by an ill-defined network of sterile inflammatory responses. The delineation of innate and adaptive immune milieu amongst the broad leukocyte infiltrate, and the gene networks which construct these responses, are poorly described in the eye. Using photo-oxidative damage in a rodent model of subretinal inflammation, we employed a novel RNA-sequencing framework to map the global gene network signature of retinal leukocytes. This revealed a previously uncharted interplay of adaptive immunity during subretinal inflammation, including prolonged enrichment of myeloid and lymphocyte migration, antigen presentation, and the alternative arm of the complement cascade involving *Factor B*. We demonstrate *Factor B*-deficient mice are protected against macrophage infiltration and subretinal inflammation. Suppressing the drivers of retinal leukocyte proliferation, or their capacity to elicit complement responses, may help preserve retinal structure and function during sterile inflammation in diseases such as AMD.

## Introduction

Leukocyte activation and recruitment is a key process in the progression of a variety of neurodegenerative diseases, including those affecting the retina (reviewed in [1]). Microglia are resident macrophages of the central nervous system, derived in early development from yolk sac myeloid cells [2, 3]. Microglia contribute to the normal physiological state of the central nervous system through constant surveillance of the parenchyma. They control a variety of processes including phagocytosis of neurons during development [4], maintenance of healthy retinal synapses [5] and tissue repair [6]. Age-Related Macular Degeneration (AMD), is a retinal disease whose hallmark is progressive loss of photoreceptors and retinal pigment epithelium (RPE), accompanied by inflammation and influx of leukocytes, including monocyte-derived macrophages [7]. Recruited macrophages, as well as resident microglia contribute to the pathogenesis of the degenerating retina (reviewed in [8, 9]), includes AMD [10–13], retinitis pigmentosa [11, 14], retinal detachment [15, 16], glaucoma [17–19] and diabetic retinopathy [17, 20]. The long-standing assumption, derived from rodent models of brain injury, is that resident microglia are the primary source of inflammatory mediators in the brain, and that recruited leukocytes do not play a long-term role in tissue repair (reviewed in [21]). However, it is also clear from recent transcriptome studies that microglia derived from different parts of the CNS have distinct molecular and functional attributes [22].

Chemokine signalling mediates monocyte migration in several CNS disorders including AMD, multiple sclerosis, Alzheimer’s disease, and brain ischemia and trauma (reviewed in [23–26]). In the photo-oxidative damage (PD) model of focal retinal degeneration we used a broad spectrum chemokine inhibitor, and reduced sub-retinal macrophage accumulation and associated photoreceptor cell death [27], indicating the key role of chemokines in retinal degenerations. Studies of Ccl2 show that ablation of Ccl2 or the Ccr2 receptor reduces monocyte infiltration and retinal degeneration in experimental choroidal neovascularization (CNV) [28] and in PD-treated Cx3cr1−/−mice [29]. Further, we have shown that expression of Ccl2 is upregulated in Müller cells in PD [30], and that targeted knockdown of Ccl2 using siRNA reduces recruitment of microglia/macrophages, photoreceptor death and complement deposition [31]. The release of endogenous triggers like Ccl2 alerts monocytes and promotes proliferation, migration, enhanced phagocytosis, as well as secretion of cytokines, chemokines, and neurotoxins (reviewed in [32]). This process of monocyte activation can promote CNS degeneration including phagocytosis of neurons that otherwise might survive [33, 34] and initiation of pro-apoptotic events [35]. The detrimental effects of macrophage aggregation in the outer retina have been directly implicated in retinal models of neovascular-AMD [36], PD [37, 38], diabetic retinopathy [39, 40], and glaucoma [41].

Despite the growing understanding of the role of macrophages in retinal degenerations, little is known about the molecular profile of these cells, nor of the contribution that other leukocytes types may have in progression of retinal disease. While macrophages are the predominant inflammatory cell in the retina, a growing body of evidence teases a broader contribution of immune responses in AMD (reviewed in [42]). Here, we report an analysis of gene expression in CD45^+^ leukocytes isolated from retina following experimental PD in a rodent model. Our constructed functional networks reveal diverse transcriptional regulation of genes involved in leukocyte activation, chemokine signalling and the complement cascade in retinal degenerations. We propose that the initial inflammatory response which primarily involves the retinal microglia/macrophages, leads to a progression in inflammatory activity, including the activation of other leukocyte subsets, initiation of an adaptive immune response, and complement activation, which contribute to subretinal inflammation and progressive photoreceptor cell death.

## Methods

### Animals

All work was conducted using either young adult Sprague-Dawley (SD) albino rats, or C57/Bl6J mice. Additionally, Cfb−/− mice (B6;129-Cfb^tm1Hrc^/Apb) were also obtained from the Australian Phonemics Facility (APF), and which were bred on the B6 background. All animal experimentation was conducted in accordance with the ARVO Statement for the Use of Animals in Ophthalmic and Vision Research and with the approval of the Animal Ethics Committee at the Australian National University, Canberra (Protocols - A2012/07 and A2014/56). Animals were raised and experiments were conducted in cyclic 5 lux light:dark (12hrs:12hrs), unless otherwise stated. All animals were culled using an overdose (60mg/kg bodyweight) of barbiturate (Valabarb; Virbac, Australia) given as intraperitoneal injection. All culling was performed at 9am to control for possible circadian effects.

### Photo-oxidative damage model

For the rat PD model, animals, housed and exposed to bright (1000 lux) light for 24hrs in accordance with our previous protocols. Exposure began and ended at 9am on successive days. Rats were euthanized for tissue collection either immediately following PD (0 days), or after a further 3 or 7 days in cyclic dim light. Dim-reared animals were collected as non-light exposed controls for comparison.

The mouse PD model was performed following our previously established methodology [43]. In brief, age-matched wild type (C57BL/6) and complement knockout animals were housed in Perspex boxes coated with a reflective interior, and exposed to 100K lux of natural white LED for up to 7 days, with access to food and water ad libitum. Each animal was administered with pupil dilator eye drops (0.1% atropine sulphate, Bausch and Lomb, Australia) two times a day during light exposure. Animals were either euthanized or subjected to electroretinogram (ERG) recordings after 3, 5, 7 days of photo-oxidative damage.

### Animal tissue collection and processing for histology and RNA extraction

Eyes from some animals were marked at the superior surface for orientation, then enucleated and immediately immersion-fixed in 4% paraformaldehyde in 0.1 M PBS (pH 7.3) for 3hrs at room temperature, then processed for cryosectioning as previously described for histological analysis [44]. From other animals (n≥6), retinas were excised through a corneal incision and prepared for cell sorting and RNA extraction.

### Analysis of Cell Death

Terminal deoxynucleotidyl transferase dUTP nick end labeling (TUNEL) was used to quantify photoreceptor apoptosis in cryosections for each experimental group, using a previously published protocol [45]. Counts of TUNEL-positive cells in the outer nuclear layer (ONL) were carried out along the full length of retinal sections cut in the parasagittal plane (superior-inferior), within the vertical meridian. The total count from each retina is the average of four sections at comparable locations.

### Outer nuclear layer (ONL) thickness measurements

Thickness of the ONL in each experimental group was measured in increments of 1mm along the full length of retinal cryosections cut in the parasagittal plane (superior-inferior), which were in close proximity to the vertical meridian. The DNA-specific dye bisbenzimide (Sigma-Aldrich, MO, USA) was used to visualize the cellular layers. The ONL thickness ratio was calculated as the thickness of the ONL relative to the distance between the ganglion cell layer (GCL) and ONL, to take into account any obliquely cut sections. The ONL thickness ratio for each retina is the average of two retinal sections at comparable locations.

### Immunohistochemistry

To detect and localise CD45^+^ cells in retinal cryosections, immunohistochemistry was performed using a CD45 primary antibody (Biolegend, San Diego, CA, USA) or IBA1 (1:500, Wako, #019-19741), as described in our previous protocols with minor modifications [46, 47]. Sections were counterstained with a DNA label (Bisbenzimide; Sigma-Aldrich) for visualisation of the retinal layers. Fluorescence in retinal sections was visualised under a laser-scanning A1^+^ confocal microscope (Nikon, Tokyo, Japan), and images were acquired using the NIS-Elements AR software (Nikon). Images were processed using Photoshop CS6 software (Adobe Systems, CA, USA).

### Flow cytometry for retinal leukocyte sub-populations

Retinal cell and dissociation and flow cytometry were conducted as per our previous protocol [43], with some modifications. Retinas from each animal were pooled and immediately placed dissociation cocktail that included 0.2% papain, and were then digested until a single cell suspension was obtained. Permeabilization was not conducted. The samples were then incubated in FC block, followed by an incubation antibody staining buffer for 45 minutes at 4°C, which contained the following cocktail of markers: CD11b-PE, 1:200, Biolegend; CD45-Alexa-647, 1:200, Biolegend; CD3-Pacific Blue, 1:400, Biolegend; CD45RA-FITC, 1:200, Biolegend; Gr1-PE, 1:200, BD Biosciences, Franklin Lakes, NJ. Samples were run through a BD Fortessa flow cytometer (BD Biosciences). The data were analysed using FloJo software (version 10.4.1). Statistical analysis was performed using Prism 6 (GraphPad Software, CA,USA). Unless specifically stated either a two-way ANOVA with Tukey’s multiple comparison post-test or an unpaired Student *t* test was utilised to determine the statistical outcome, with a *P* value of <0.05 considered statistically significant.

### Fluorescence-activated cell sorting of CD45+ Leukocytes

Retinas from each animal were pooled and immediately placed in chilled HBSS (n = 3 per time point) and then subjected to light mechanical separation using a razor blade. Samples were transferred into 0.2% papain digestion cocktail [47] and incubated at 8°C for 45 minutes, then 28°C for 7 minutes. The resulting homogenate was centrifuged at 250 g for 5 minutes at 4°C, and the pellet was resuspended in neutralization buffer. The homogenate was centrifuged again at 420g for 5 minutes at 4°C, and the pellet resuspended in staining buffer containing 1.0% bovine serum albumin (BSA), and 0.1% azide. The samples were incubated in staining buffer containing anti-CD45-Alexa 647 (Biolegend) for 45 minutes at 4°C, then washed twice in HBBS and resuspended in staining buffer. The resultant CD45-stained samples were run through a fluorescence-activated cell sorter (FACS) (BD FACSAria II; BD Biosciences, Franklin Lakes, NJ, USA). Viability of the sorted cells was assessed by labeling with DAPI. The isolated CD45+ cells were collected in staining buffer and kept chilled on ice until RNA extraction could be commenced. To prepare for RNA extraction, isolated samples were centrifuged at 420 g for 5 minutes at 4°C, and the supernatant removed. RNA extraction was performed with a combination of TRIzol Reagent (Life Technologies,Carlsbad, CA, USA) and an RNAqueous-small scale kit (Life Technologies, Carlsbad, CA, USA) utilized in tandem to extract and purify the RNA respectively. Isolated total RNA was analysed for quantity and purity with a ND-1000 spectrophotometer (Nanodrop Technologies, Wilmington, DE, USA).

### RNA-seq sample preparation

RNA samples were prepared for RNA-seq by the Australian Cancer Research Foundation (ACRF) Biomolecular Resource Facility (BRF) (John Curtin School of Medical Research, Australian National University, Canberra, Australia). cDNA library preparation was performed using the SMARTer Low RNA Kit (Clontech, Mountain View, CA, USA) as per the manufacturer’s instructions, using 679pg of RNA per sample was used as starting material for the preparation of the cDNA libraries. Following library preparation, DNA was simultaneously fragmented and tagged with sequencing adaptors using an adapted Nextera DNA Sample preparation protocol (Illumina Technologies, San Diego, CA, USA – Revision A), using Ampure beads for purification (Agilent Technologies, Santa Clara, CA, USA). Concentrations of libraries were checked on the Agilent Bioanalyzer (Agilent Technologies) and pooled to equimolar amounts. Fragmented samples were run on the HiSeq2500 (Illumina Technologies), using TruSeq^®^ Rapid PE Cluster Kit 2500 (Illumina Technologies) and TruSeq^®^ Rapid SBS Kit 2500 – 200 cycles – (Illumina Technologies). 100bp paired end sequencing was performed on all samples (12 sample, 4 data points in triplicate – 18.33 million reads per sample).

### RNA-seq alignment and analysis

All sequenced data was assessed for quality using FastQC software [48] followed by filtering of low quality data using Trimmomatic software (version 0.27, LEADING:15 TRAILING: 15 SLIDINGWINDOW:4:20 MINLEN:60 trimming parameters) [49]. All sequence data remaining as paired data after quality filter were mapped to the Rat genome (assembly version Rnor_5.0) using TopHat software (v2.0.10, --b2-very-sensitive mapping parameter) [50] against gene annotations obtained from the Ensembl v75 database. Samtools (v0.1.19.0) was then used to remove unmapped reads and secondary alignments from TopHat output (fixmate −r parameters) [51]. HTSeq was used for obtaining tag counts for each annotated gene using default parameters for non-strand specific library [52]. Tag counts were normalized for library composition and library size using Trimmed Mean of M-values (TMM) method as implemented in edgeR package to obtain counts per million (CPM) [53, 54] Sum of tag counts in each sample was used as the effective library. All RNAseq data for this project is available on NCBI Short Read Archive with the BioProject ID SRP133267.

Differential expression analysis compared the normalised CPM values of the light-treated groups to the control cohort (dim), over a time course of 0, 3, and 7 days in a pair-wise manner. Genes were considered differentially expressed if they had a p-value if < 0.05 and false discovery rate (FDR) of < 0.05 (One-way ANOVA); a fold change cut-off of 1.5 was also applied. The relatedness of the individual samples was assessed with principal component analysis (PCA) on the log2 cpm of the DEGs derived from each comparison, and which utilised the scatterplot3d package in R (v3.2.2). Venn-diagrams were also created to illustrate overlap between data points in these gene sets (Venny, v2.1). Gene co-regulation across the time course was assessed by K-means clustering analysis on the log2 cpm of the DEGs using the Stats Bioconductor package in R (v2.15.0) (scripting available upon request). The data were further examined with heatmaps using hierarchical clustering via Euclidian distance, which was conducted with heatmapper [55].

Gene ontology (GO) term enrichment analysis was performed using the online bioinformatics resource Panther (v13.0) to identify overrepresented biological processes at each time point. GO analysis was conducted using statistical overrepresentation, with a Bonferroni correction applied to account for multiple testing. Additionally, pathway analysis was performed using the Reactome online database (v62) to interrogate the data for statistically enriched pathways, as ranked by FDR (<0.05). To improve the interpretation of the GO terms obtained from Panther, an integrated network analysis was employed using ClueGO (v2.3.3), a plug-in for Cytoscape Software (v3.5.1) [56], using DEGs from each comparison. Networks were constructed using GO terms for biological process, and used enrichment/depletion hyper-geometric distribution tests with an adjusted p-value of <0.05 (Bonferroni) for terms and groups. Kappa-statistics score threshold was set to 0.5 to define the functional grouping, while the leading term for groups was selected based on highest significance. GO term fusion was applied to reduce the redundancy of the terms included in the networks.

### Electroretinogram recordings

Full-field scotopic ERG recording assessed the retinal function between CfB−/− and Wt mice, and used our previously published methodology [43]. Briefly, a flash stimuli for mixed responses were provided by an LED-based system (FS-250A Enhanced Ganzfeld, Photometric Solutions International, Melbourne), over a stimulus intensity range of 6.3 log cdsm-2 (range-4.4-1.9 log cdsm-2). The a-wave amplitude was measured from the baseline to the trough of the a-wave response and the b-wave amplitude was measured from the trough of the a-wave to the peak of the b-wave. Data are expressed as the mean response amplitude ± SEM (μV). Two-way ANOVA, with Tukey’s multiple comparisons Post-hoc test, was performed to compare the responses over the flash stimulus range.

### Western blotting for C3d

Whole retinas from euthanized Wt or C3b−/− mice were collected at 4°C in CellLytic M buffer (Sigma, Australia) containing protease inhibitor cocktail (Roche), and spun down at 13,000g to obtain the extract protein. Concentration of samples was determined by Bradford assay (Bio-Rad), and equal amounts of total proteins were loaded onto 4-20% Mini-PROTEAN Tris-Glycine gels (Bio-Rad). Following electrophoresis (200V for 35 min), proteins were transferred to nitrocellulose membranes (Bio-Rad) in semi-dry transfer system for 30 min −2 hr depending on the size of protein. Blots were blocked with 3% BSA, 0.01% Tween 20, and probed overnight at 4°C with antibodies for either C3d (1:500, #AF2655-SP, R&D Systems) or the loading control GAPDH (1:4000, #G9545, Sigma Aldrich). Immunoblots were incubated with HRP-conjugated secondary antibodies for 2 hr at room temperature and developed using Clarity ECL Western Blotting Substrate (Bio-Rad). Visualization and imaging of blots was performed on ChemiDoc MP Imaging System (Bio-Rad).

## Results

### Photoreceptor cell apoptosis and degeneration

Exposure to PD led to an increase in TUNEL-positive cells in the ONL (**Figure 1A**), and a subsequent decrease in ONL thickness (**Figure 1B**) compared to dim-reared controls. At 7 days post-exposure, persistent ONL thinning and photoreceptor cell death was observed, consistent with photoreceptor death, as described previously [57]. Photoreceptor cell death was correlated with both increased numbers and altered distribution of CD45^+^ leukocytes in the retina (**Figure 2A-D**). CD45+ cells in retinas of dim-reared control animals showed localisation of ramified leukocytes predominantly in the inner retina, with some cells present in the choriocapillaris (**Figure 2A**). At 7 days post-PD, CD45^+^ cells had an activated/rounded in morphology, and were distributed throughout all layers of the retina including the ONL and the subretinal space (**Figure 2B-D**). Quantification of total retinal leukocytes by flow cytometry showed increased numbers of CD45^+^ cells following PD, with highest numbers detected at 7 days (**Figure 2E**). In dim-reared control animals the percentage of CD45^+^ cells in the total retinal population was 0.85% compared with to 6.2% 7 days post-damage (P<0.05, ANOVA).

**Figure 1.**
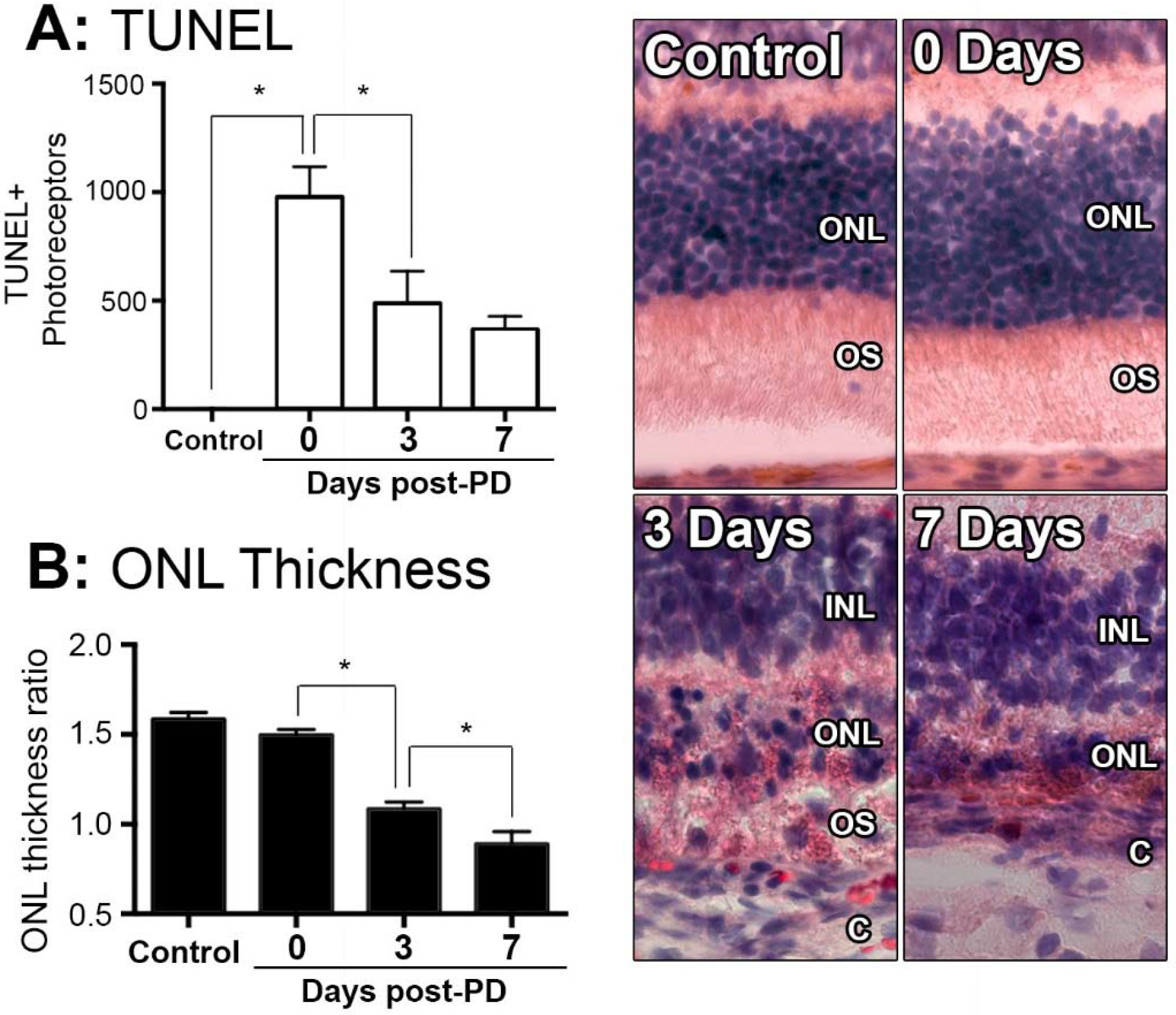
Changes in photoreceptor apoptosis and degeneration following photo-oxidative damage. **A:** TUNEL+ photoreceptors were quantified across retinal sections for each replicate in each group. There was immediately increase in TUNEL+ photoreceptors following photo-oxidative damage, at 0 days, which remained significantly elevated at 3 and 7 days post-exposure (P<0.05). **B:** ONL thickness was averaged across retinal sections and expressed as a ratio of the total retinal thickness, for each sample. There was a progressive decrease in ONL thickness from 0 days to 7 days post-exposure (P<0.05). Representative images are taken from the superior retina, approximately 500μm from the optic nerve head. The data depict a sample size of n=5 per group. Asterisks equates to p-value of <0.05, as determined by Tukey’s post-hoc test. ONL, outer nuclear layer; OS, outer segments; INL, inner nuclear layer; C, choroid.

### Temporal profiling of retinal CD45+ leukocytes and subpopulations

Common lineage markers for macrophages (CD11b), granulocytes (Gr1), T cells (CD3), and B cells (CD45RA) were used to investigate the major subsets of CD45+ leukocytes in the retina over the damage time course (**Figure 2F-H**). The CD11b+ macrophages were found to increase relative to the retinal population over the experimental time course. This peaked at 7 days post-exposure (P<0.05, ANOVA), and detected on 5.1% of the retinal cell population (**Figure 2F**). In control samples, 57% of CD45^+^ cells positive for CD11b, while at 7 days 85% of the CD45+ cells were also CD11b+, indicating that changes in the CD45+ population broadly reflects the changes present in the CD11b+ population. The subset of GR1+CD11b+ granulocytes, broadly encompassing neutrophils and eosinophils, were shown to increase from near zero in dim-reared controls to ~1.5% of the retinal population at 7 days post-damage (P<0.05 ANOVA) (**Figure 2G**). The subset of CD3+ T cells displayed progressive increase over time, and by 7 days comprised 20% of the CD45+CD11b-population, which was almost quadruple the proportion in the dim-reared controls (P<0.05 ANOVA) (**Figure 2H**). CD45RA+ B cells in contrast were barely dateable and showed no appreciable change across the PD time course (**Figure 2H**).

**Figure 2.**
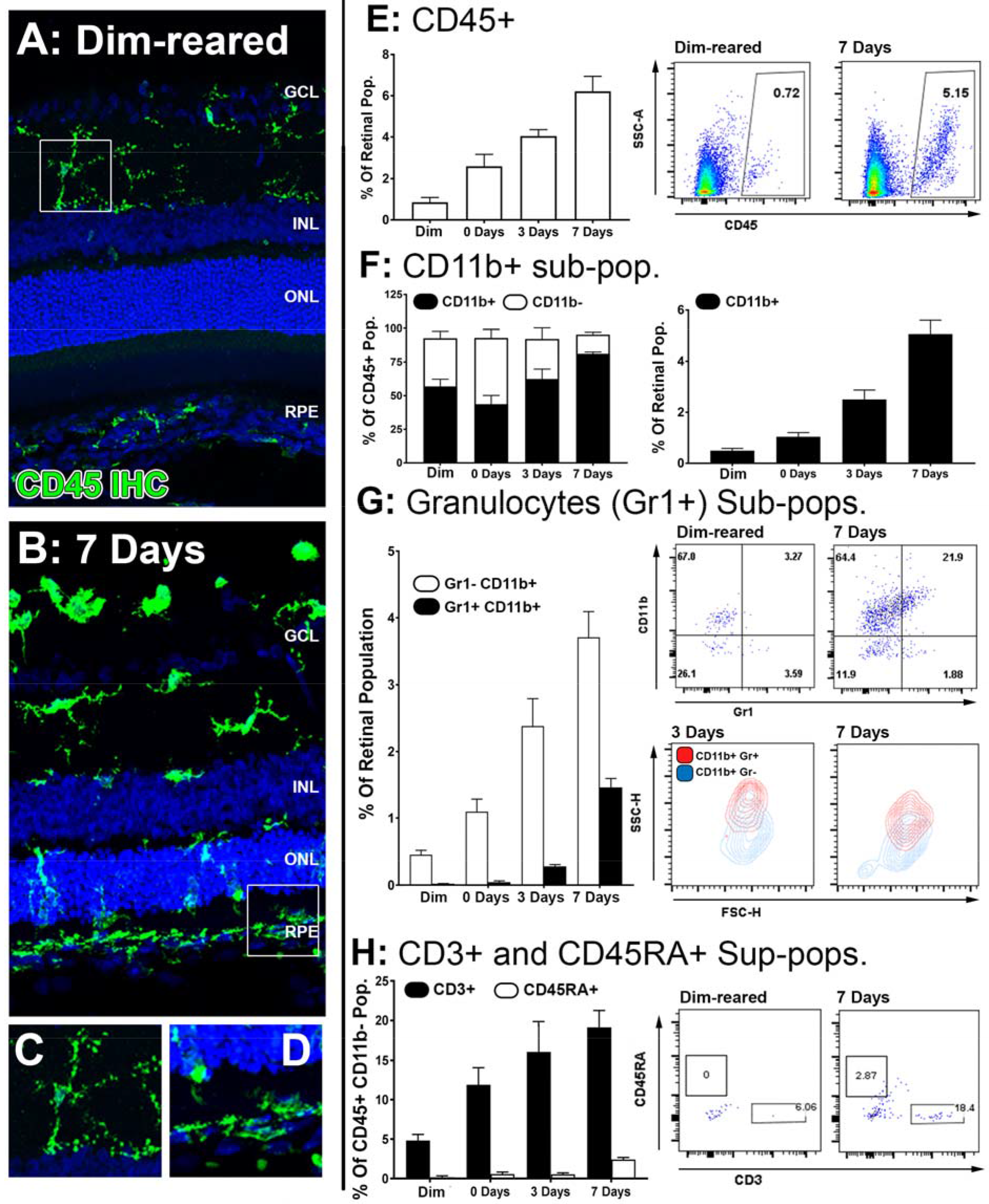
Characterisation of major CD45 leukocyte subsets in the retina following photo-oxidative damage. **A-D:**Representative labelling of CD45+ cells (green) by immunohistochemistry on retinal sections, with DAPI (blue) as the background stain. In dim-reared controls, staining for CD45 was restricted to ramified microglia in the inner plexiform layer IPL (A,C). By 7 days post damage, there were numerous CD45+ cells distributed in the ONL and subretinal space following damage (B,D). **E:** Graphing and representative plots for florescence activated cell sorting (FACS) of CD45+ cells, which show progressive increases over time, relative to the retinal population (P<0.05, ANOVA). **F-H:** Categorisation of CD45 subpopulations in the retina over the timecourse, using flow cytometry. **F:** The proportion of CD11b+ macrophages comprised the bulk of the CD45+ population (F, left, P<0.05, ANOVA), and continually increased throughout the timecourse relative to the retinal population (F, right, P<0.05, ANOVA). **G:** Graphing and representative plots showcase CD11b and Gr1 staining amongst CD45 cells, relative to the retinal population. CD11b+Gr1+ granulocytes were found exhibit late increase at 3 and 7 days (ANOVA P<0.05), comprising a smaller proportion than the CD11b+Gr1- macrophage subset. The representative SSC/FSC plots at 3 and 7 days show increased SSC of CD11b+Gr1+ Granulocytes compared to CD11b+Gr1-. **H:** Staining for CD3+ T cells and C45RA+ B cells is presented in graphs and representative plots as a proportion of the CD45+CD11b- parent population. The subset of CD3+ cells displayed progressive increase over time, and peaking at 7 days (P<0.05 ANOVA); CD45RA+ cells in contrast showed no appreciable change across the timecourse. All datasets represent a sample size of n=4 per group. C, choroid; FACS fluorescence activated cell sorting; INL, inner nuclear layer; ONL, outer nuclear layer; OS, outer segments.

### Transcriptome profile of isolated CD45^+^ retinal leukocytes

RNAseq was performed on CD45+ cells isolated from the retinas over 0, 3 and 7 days post-PD to construct a temporal transcriptional fingerprint of immune modulation. PCA was performed on the individual samples for all expressed genes (**Figure 3**). The scree plot demonstrated that most of the variance was observed in the first 3 principle components (**Figure 3A**). The PCA indicated a strong correlation between sample replicates, and highlights a rapid divergence in transcriptional profile between dim-reared control leukocytes, and those isolated at the point of initial injury at 0 days (Figure 3B). Though distinct, the 3 and 7 day groups appeared more closely clustered together than 0 days, and the 7-day grouping appeared the most similar to the control cluster (**Figure 3B**).

**Figure 3.**
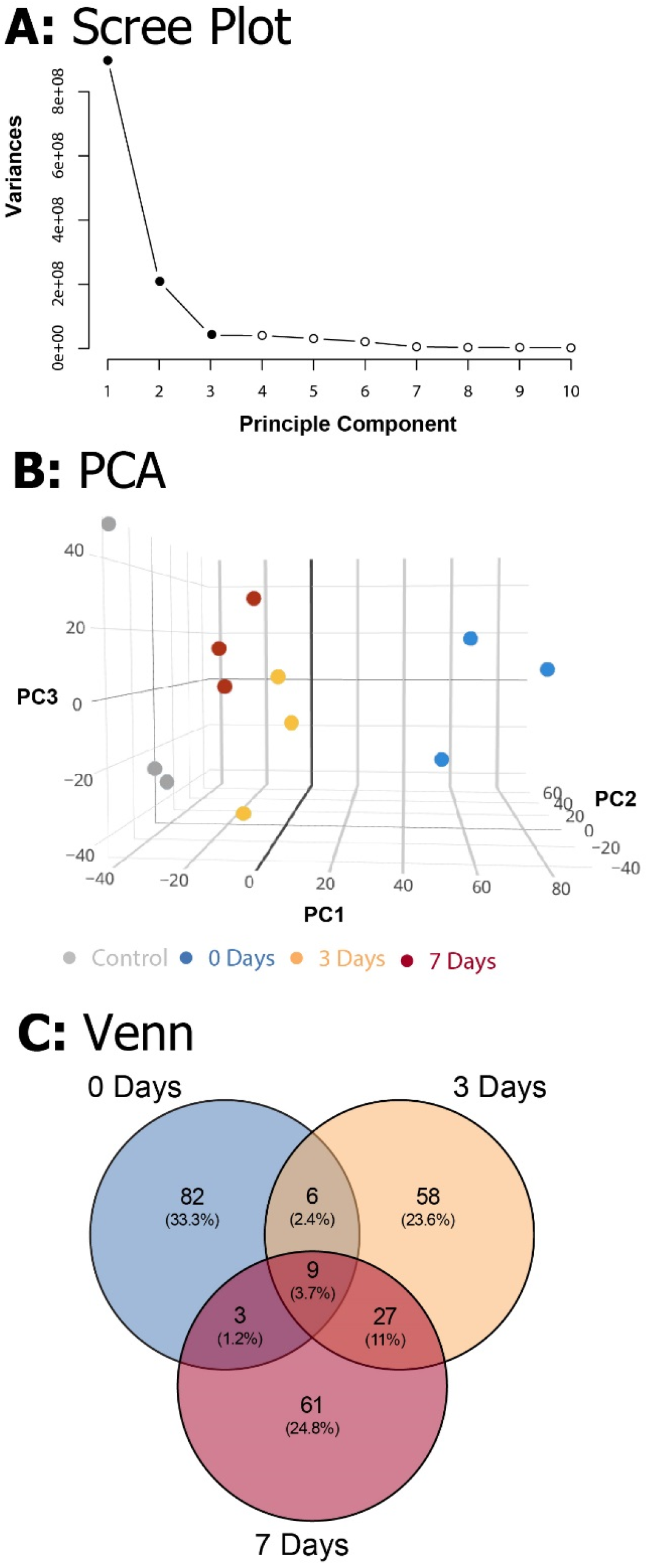
Principle Component Analysis of the RNAseq dataset of isolated CD45+ retinal leukocytes. PCA was conducted on the log2cpm of all expressed genes. **A:** The scree plot showcases the variance across 10 PCs; the vast majority of the variance is explained in first 3 PCs (highlighted in black). **B:** PCA graphed the PCs 1-3 for all sample groups in triplicate, and indicated distinct clustering for replicates among respective groups. **C:** Venn diagram depicts the distribution and interrelation of the top 100 DEGs at each of the post-damage time points. DEG, differentially expressed gene; PC, principle component; PCA, principle component analysis.

When compared to control samples, 2193 genes were found to be significantly differentially expressed in the light-treated groups (adjusted p<0.05, **Supplementary Table S1**) with 1818 DEGs identified at 0 days, and progressive decline to 656 genes at 3 days, and 155 genes at 7 days. As this variance in number of DEGs has the potential to heavily bias downstream comparisons, we took the top 100 most significant DEGs for each time point, as illustrated in the volcano plots and PCA in **Supplementary Figure S1** to take forward for comparative functional enrichment analysis. These were representative of the trend in both Venn and PCA analyses that were drawn from entire DEG list (**Supplementary Figure S2**). Of the combined total of 300 DEGs, relatively few genes were identified in more than one time point, resulting in a final representative snapshot of 246 unique DEGs (**Figure 3C**). Of these, 82 (33.3%) were found exclusively at 0 days, compared to 58 (23.6%) and 61 (24.8%) for 3 and 7 day groups respectively. The 3 and 7 day groups were also found to share the most DEGs (27, 11%), as opposed to a total of only 9 (3.7%) with the 0 days group. These patterns indicate the greatest transcriptional changes in retinal leukocytes occurring within the first 24 hours, and most of these are acutely resolved within a week after injury.

### Network analysis of CD45+ leukocyte transcriptome

To gain insight into the biological processes (BP) which mark the temporal shift in the CD45+ transcriptome over the course of pathology, GO analysis was performed on the list of Top 100 DEGs at each point (**Supplementary Table S2**). Lists of GO:BP terms to showcase the Top 10 terms ranked by significance for each point (**Table 1**, adj. p<0.05). A striking observation from this analysis, however, was the extent to which these enriched biological processes were being driven by small clusters of genes (**Table 1**, network genes). This was particularly evident at the 0 day point, wherein a subset of chemokines (*Ccl2*, *Cxcl16*, *Cxcl11*, *Ccl12*, *Ccl7*, *Ccl22*, and *Ccl17*) was shown to underscore almost all the biology illustrated by the top ranked terms.

**Table 1.** Functional Overrepresentation of GO:BP Terms in Panther

| GO biological process complete | Background Set | Network Set | Network Fold Enrichment | Adjusted p-value | Network Genes |
| --- | --- | --- | --- | --- | --- |
| Control vs 0 Days |  |  |  |  |  |
| lymphocyte chemotaxis (GO:0048247) | 34 | 8 | 69.26 | 5.14E-09 | Adam8, Ccl2, Cxcl16, Cxcl11, Ccl12, Ccl7, Ccl22, Ccl17 |
| T cell migration (GO:0072678) | 19 | 4 | 61.97 | 6.73E-03 | Ccl2, Cxcl16, Cxcl11, Itgb7 |
| lymphocyte migration (GO:0072676) | 45 | 9 | 58.87 | 7.12E-10 | Adam8, Ccl2, Cxcl16, Cxcl11, Ccl12, Ccl7, Itgb7, Ccl22, Ccl17 |
| monocyte chemotaxis (GO:0002548) | 30 | 6 | 58.87 | 1.24E-05 | Anxa1, Ccl2, Ccl12, Ccl7, Ccl22, Ccl17 |
| mononuclear cell migration (GO:0071674) | 33 | 6 | 53.52 | 2.18E-05 | Anxa1, Ccl2, Ccl12, Ccl7, Ccl22, Ccl17 |
| granulocyte chemotaxis (GO:0071621) | 65 | 7 | 31.7 | 3.38E-05 | Anxa1, Ccl2, Spp1, Ccl12, Ccl7, Ccl22, Ccl17 |
| neutrophil chemotaxis (GO:0030593) | 58 | 6 | 30.45 | 6.00E-04 | Ccl2, Spp1, Ccl12, Ccl7, Ccl22, Ccl17 |
| granulocyte migration (GO:0097530) | 71 | 7 | 29.02 | 6.17E-05 | Anxa1, Ccl2, Spp1, Ccl12, Ccl7, Ccl22, Ccl17 |
| chemokine-mediated signaling pathway (GO:0070098) | 61 | 6 | 28.95 | 8.05E-04 | Ccl2, Cxcl11, Ccl12, Ccl7, Ccl22, Ccl17 |
| neutrophil migration (GO:1990266) | 63 | 6 | 28.03 | 9.72E-04 | Ccl2, Spp1, Ccl12, Ccl7, Ccl22, Ccl17 |
| Control vs 3 Days |  |  |  |  |  |
| regulation of chromosome segregation (GO:0051983) | 86 | 6 | 19.99 | 6.93E-03 | Ube2c, Dusp1, Pttg1, Knstrn, Bub1b, Spag |
| mitotic sister chromatid segregation (GO:0000070) | 95 | 6 | 18.1 | 1.23E-02 | Aurk, Pttg1, Knstrn, Kif22, Cxc20, Spag5 |
| mitotic nuclear division (GO:0140014) | 131 | 8 | 17.5 | 2.38E-04 | Ube2c, Aurkb, Pttg1, Knstrn, Tpx2, Kif22, Cdc20, Spag5 |
| sister chromatid segregation (GO:0000819) | 119 | 7 | 16.86 | 2.42E-03 | Ube2c, Pttg1, Knstrn, Bub1b, Kif22, Cdc20, Spag5 |
| spindle organization (GO:0007051) | 122 | 7 | 16.44 | 2.86E-03 | Ube2c, Cep72, Knstrn, Stmn1, Tpx2, Cdc20, Spag5 |
| nuclear division (GO:0000280) | 263 | 11 | 11.99 | 2.35E-05 | Ube2c, Aurkb, Rad51, Pttg1, Knstrn, Bub1b, Tpx2, Kif22, Cdc20, Spag5 |
| regulation of nuclear division (GO:0051783) | 170 | 7 | 11.8 | 2.55E-02 | Ube2c, Tnf, Dusp1, Pttg1, Bub1b, Cdc20, Ifg1 |
| mitotic cell cycle process (GO:1903047) | 395 | 16 | 11.61 | 5.61E-09 | Ube2c, Aurkb, Iqgap3, Rad51, Kif20a, Pttg1, Knstrn, Bub1b, Stmn1, Cdkn3, Tpx2, Kif22, Ier3, Cdc20, Spag5 |
| mitotic cell cycle (GO:0000278) | 457 | 18 | 11.29 | 2.58E-10 | Ube2c, Aurkb, Iqgap3, Nuf1, Rad51, Kif20a, Pttg1, Cenpw, Knstrn, Bub1b, Stmn1, Cdkn3, Tpx2, Kif22, Ier3, Cdc20, Spag5 |
| organelle fission (GO:0048285) | 294 | 11 | 10.72 | 7.35E-05 | Ube2c, Aurkb, Rad51, Pttg1, Knstrn, Bub1b, Tpx2, Kif22, Cdc20, Spag5 |
| Control vs 7 Days |  |  |  |  |  |
| antigen processing and presentation of peptide antigen via MHC class I (GO:0002474) | 29 | 4 | 46.22 | 2.11E-02 | RT1-T24-4, RT1-CE10, RT1-CE5, RT1-CE5 |
| antigen processing and presentation of peptide antigen (GO:0048002) | 47 | 6 | 42.78 | 7.93E-05 | RT1-T24-4, RT1-CE10, RT1-CE5, Cd74, RT1-CE5, RT1-Da |
| antigen processing and presentation (GO:0019882) | 79 | 6 | 25.45 | 1.66E-03 | RT1-T24-4, RT1-CE10, RT1-CE5, Cd74, RT1-CE5, RT1-Da |
| cell adhesion (GO:0007155) | 654 | 11 | 5.64 | 3.79E-02 | Itgae, Tnf, Gpnmb, Axl, Rom1, Cd63, Mfge8, Cldn1, Lgals3bp, Cdh17, Cd96 |
| biological adhesion (GO:0022610) | 663 | 11 | 5.56 | 4.32E-02 | Itgae, Tnf, Gpnmb, Axl, Rom1, Cd63, Mfge8, Cldn1, Lgals3bp, Cdh17, Cd96 |
| immune response (GO:0006955) | 798 | 13 | 5.46 | 6.23E-03 | Fut7, Tnf, RT1-T24-4, Cfb, Cxcl13, Cldn1, RT1-CE5, Cd74, Cdh17, RT1-CE5, Skap1, RT1-Da |
| cell surface receptor signaling pathway (GO:0007166) | 1591 | 18 | 3.79 | 6.48E-03 | Itgae, Tnf, Card14, Axl, Csf2ra, Sulf1, Evc, Rom1, Cd63, Adgre5, Cxcl13, PLxnc1, Cxcr6, Cxcr3, Cd74, Cdh17, Cd3e, Skap1 |
| immune system process (GO:0002376) | 1607 | 18 | 3.75 | 7.50E-03 | Ctnnb1, Fut7, Tnf, Axl, RT1-T24-4, Cfb, Ifitm1, Cxcl13, Cldn1, RT1-CE10, Cxcr3, RT1-CE5, Cd74, Cdh17, RT1-CE5 Cd3e, Skap1, RT1-Da |

To facilitate a better understanding of the biological significance of the data, we used ClueGO to integrate enriched GO:BP terms from each point and organise them into a combined functional network (**Figure 4**). This enabled us to explore functional interrelationships across a broad biological network, whilst also understanding redundancy of terms between related functions. **Figure 4A** represents the global functional network of retinal CD45+ population using GO:BP terms constructed from the combined Top 100 DEGs from each time point, and showcase functional grouped clusters such as lymphocyte migration, positive regulation of hemopoiesis, and nuclear division. In **Figure 4B**, enrichment of specific points across the network is displayed as proportion (%) of associated genes (complete readout of GO terms and gene are tabulated in **Supplementary Table S3**). Here, the data reveal a shift in enrichment over the course of PD. The 0 days point was most represented in the lymphocyte migration functional group (69.2%), as well as networked terms for monocyte and neutrophil migration (61.4, 66.7%), extravasation (100%) interleukin 1 (80%), conversely, there was pronounced enrichment of functional groups that underscore cellular proliferation and metabolism, including nuclear division (89.4%) and glycolytic process (68.3%). After 7 days, there was a shift toward adaptive immune response, with pronounced representation of terms relating to antigen processing and presentation (74%), including MHC class I (100%), and T cell receptor signalling pathway (77.4%). Individual functional networks were also constructed for the individual time points (**Supplementary Figure S3**), though were found to explain the data in a largely similar fashion to the combined network.

**Figure 4.**
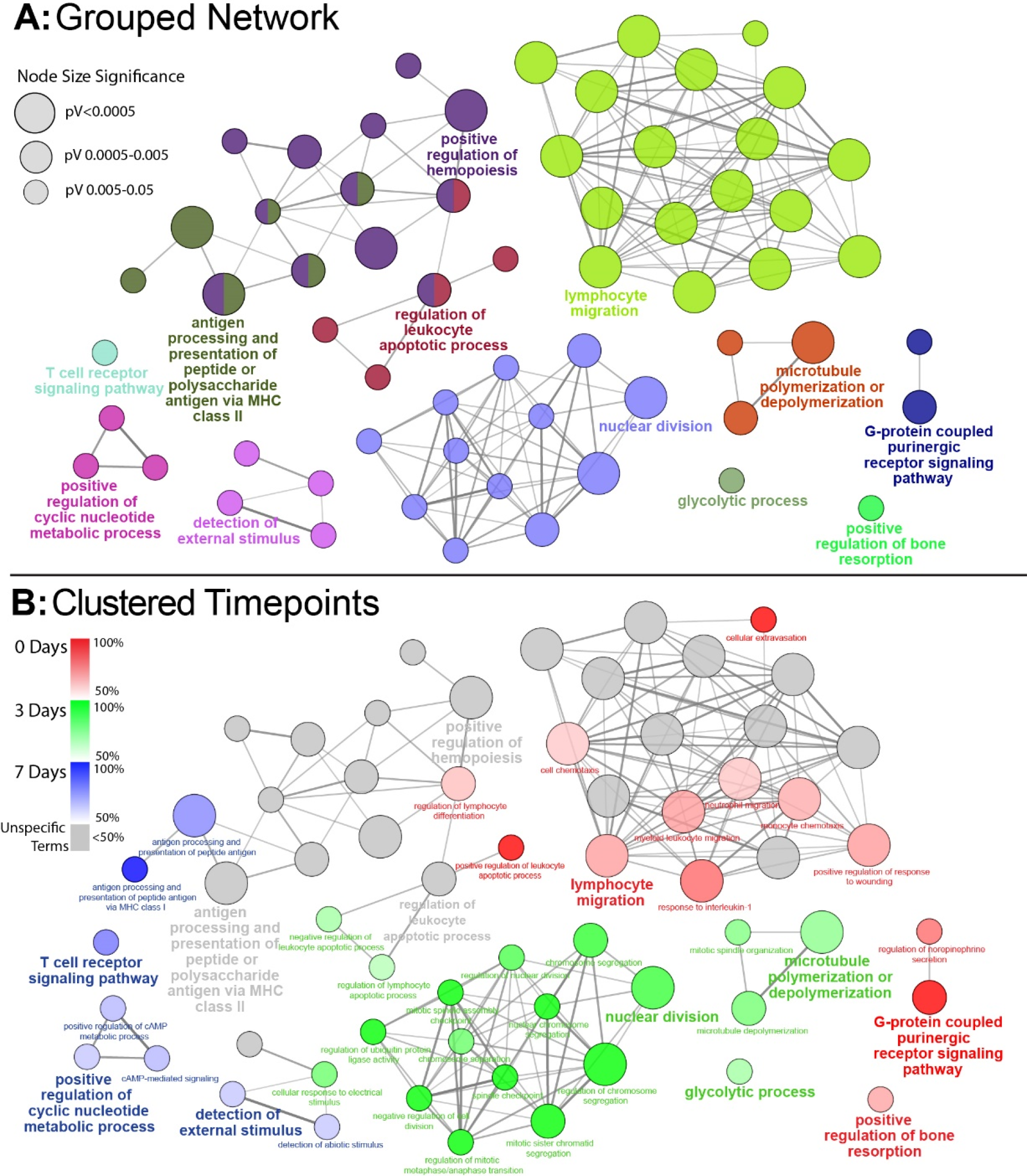
Functionally grouped network analysis of enriched GO terms for CD45 Population over the timecourse. **A:** The top 100 DEGs from each timepoint were used to generate enriched GO terms for biological process, which were then integrated into a functionally grouped network. GO:BP terms are represented as nodes, while their size illustrates the significance of the term enrichment; the edges reflect the degree of connectivity and grouping between the terms. The leading term in each functional grouping is selected based on the highest significance (Bonferroni adj. P<0.05). **B:** The network is overlayed with the enrichment of the timepoints across the functional groupings; nodes in which more than 50% of the genes are attributed to a given timepoint are colour-coded accordingly. BP, biological process; DEG, differentially expressed gene; GO, gene ontology.

### Gene co-regulation using K-means clustering

The co-regulation of DEGs over the damage time course was assessing using K-means clustering, which was performed on the combined list of 246 unique DEGs. The analysis identified 4 major clusters as shown in heat maps and graphs in **Figure 5**; more detailed heat-maps showing the specific genes in each cluster are located in **Supplementary Figure S4**. Based on their temporal expression profile, the clusters were variously classed as Early Up **(A)**, Mid Up **(B)**, Late Up **(C)**, and Global down **(D)**, and GO:BP and Reactome Pathway (RP) analysis were performed on each to identify significantly enriched terms and pathways. The Top 10 entries for each analysis (ranked by adj. P value) are listed in **Figure 5A-D** the (complete list is available in **Supplementary Table S4**).

**Figure 5.**
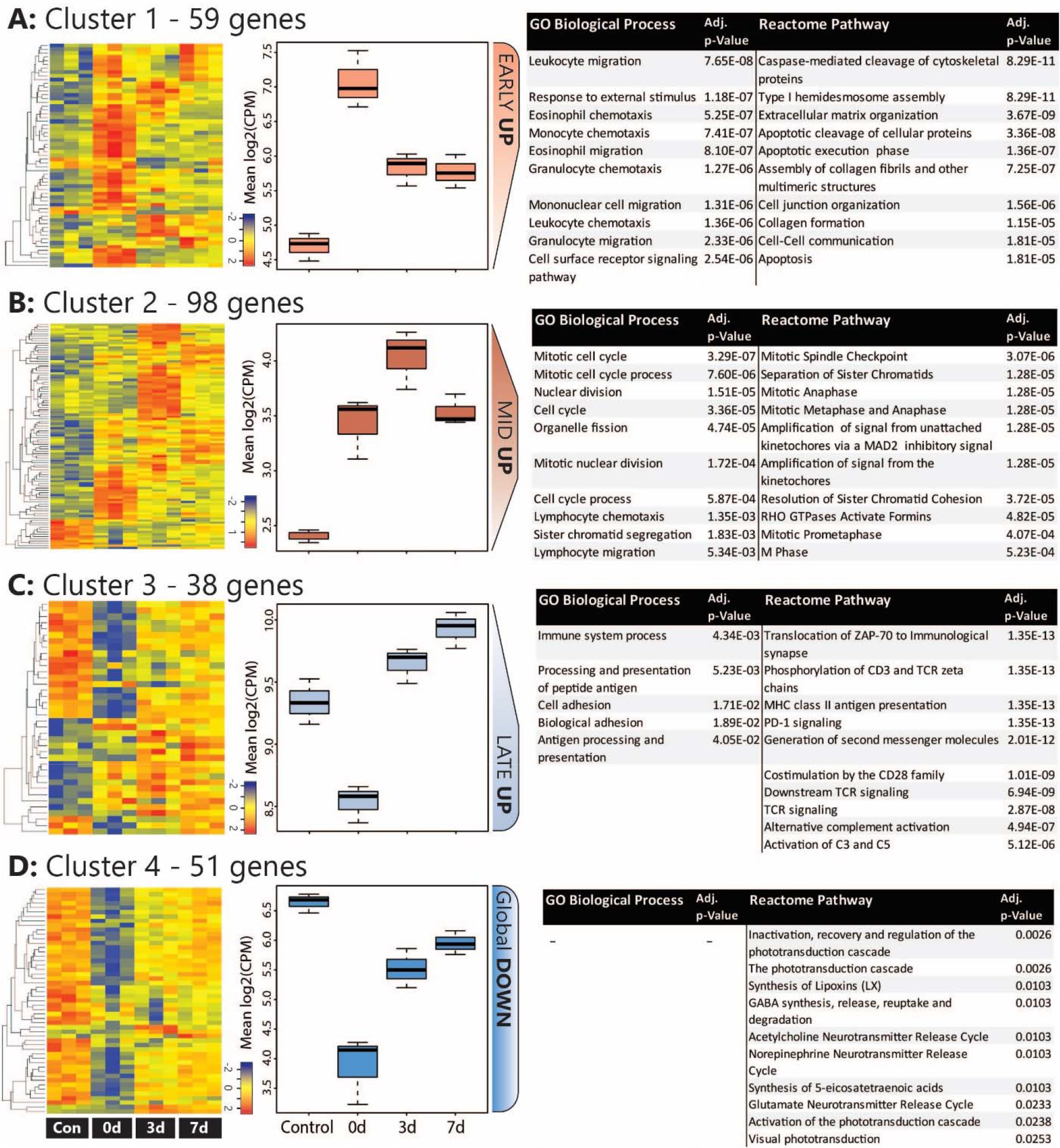
Analysis of co-regulated DEGs within the CD45 Population following photo-oxidative damage. **A-D:** K-means clustering was performed on the log2cpm of 246 DEGs that were derived from the combined Top 100 DEGs in each timepoint. 4 major clusters were identified, as illustrated in the heat maps and box and whiskers plots, and variously termed **(A)** Early UP, **(B)** Mid UP, **(C)** Late UP, and **(D)** Global DOWN, based on their temporal expression pattern. The DEGs in each cluster were assessed for enriched terms and pathways using GO:BP and Reactome, respectively; the Top entries for each are displayed, which were ranked according to the adjusted P value. BP, biological process; DEG, differentially expressed gene; GO, gene ontology.

The Early Up cluster (**Figure 5A**) identified DEGs that exhibited peak up-regulation at 0 days, and mainly comprised GO terms associated with recruitment, including leukocyte migration (GO:0050900), monocyte chemotaxis (GO:0002548), and granulocyte chemotaxis (GO:0071621). These consisted of many overlapping chemokines such as Ccl2, *Ccl3*, *Ccl7*, *Ccl12*, as well as the pro-inflammatory regulators *Anxa1* (Annexin A1) and *Spp1* (Osteopontin), and the immune suppressor *lgals1* (Galectin-1). Pathway analysis, conversely, identified many apoptosis-related entries such as caspase-mediated cleavage of cytoskeletal proteins (R-RNO-264870) and apoptotic execution phase (R-RNO-75153). These enriched chemokine pathways are representative of the entire DEG list at the 0 days timepoint (**Supplementary Figure S5**).

Mid Up (**Figure 5B**) correlated with peak up-regulation at 3 days. Both GO terms and pathways associated with this time point were dominated by proliferative responses, including mitotic cell cycle (GO:0000278) and Mitotic Spindle Checkpoint (R-RNO-69618). Also enriched, however, was lymphocyte chemotaxis (GO:0048247), consisting of the *Adam8* and the chemokines *Cxcl9*, *Cxcl11*, *Ccl22*, *and Ccl17*. Other chemokines in this cluster also include the *Xcl1-Xcr1* signalling axis (**Supplementary Figure 3A**). Late up (**Figure 5C**) clustered DEGs that exhibited peak upregulation and 7 days, and revealed terms for antigen processing and presentation of peptide antigen (GO:0048002), and pathways associated with T-cell activation, including Translocation of ZAP-70 to Immunological synapse (R-RNO-202430), Phosphorylation of CD3 and T cell receptor (TCR) zeta chains (R-RNO-202427). Strikingly, the complement activator Factor B (*cfb*) was also present in the Late-up cluster, alongside an enrichment of pathways associated with Alternative complement activation (R-RNO-173736) and Activation of C3 and C5 (R-RNO-174577); these are further illustrated in pathway diagrams depicted in **Figure 6**. The Global-DOWN Cluster (**Figure 5D**), unlike the other groupings, did not return any significantly enrich terms for GO:BP.

**Figure 6.**
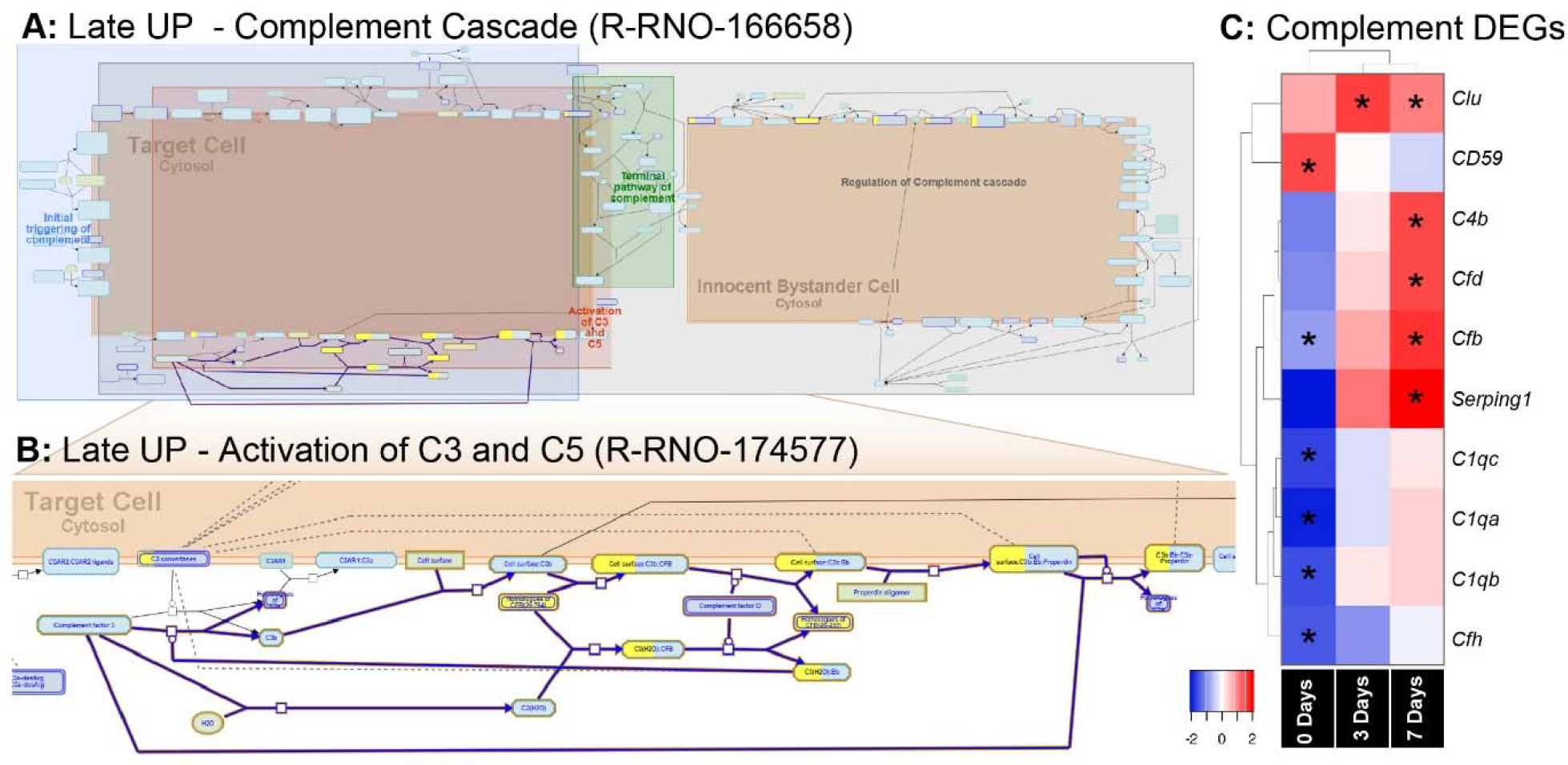
Enrichment of the complement pathway in the CD45 Transcriptome. **A-B:**Reactome pathway diagrams showcase overrepresentation (FDR<0.05) of complement interactions in the Late UP K-means Cluster (Figure 5). Yellow-colouring in the nodes denotes a gene in the expression list that matches a protein within the pathway diagram, while the degree of colouring represents the coverage. **A:** Pathway diagram depicts the entire complement cascade, which are broadly classified subcomponents for Initial triggering of complement (blue), Terminal pathway of complement (green), Activation of C3 and C5 (red), and Regulation of the complement cascade (grey). **B:** Activation of C3 and C5 was found to show significant enrichment Factor B (*cfb*) in the expression list (yellow-colouring). **C:** Heat maps with hierarchical clustering on significant complement DEGs derived from the full gene expression list (2193). Asterisks denote significant differential expression (FDR<0.05) compared to control (dim). DEG, differentially expressed gene.

### Effect of Factor B ablation on subretinal inflammation following PD

The pronounced enrichment of alternative complement pathway and Factor B within the K-means data led us to further explore its role in subretinal inflammation and photoreceptor degeneration after photo-oxidative damage. Interrogation of the full list of 2193 DEGs revealed further significant differences in the complement genes, including alternative Factor D (*Cfd*), and classical components *C1qa*, *C1qb*, and *C1qc* (**Figure 7C**). Hierarchical clustering on the genes revealed an early decrease (0 days) in the cluster of C1q components, while both the alternative pathway Factors B and D showed increased expression at 7 days, in agreement with the K-means pathway analysis.

**Figure 7.**
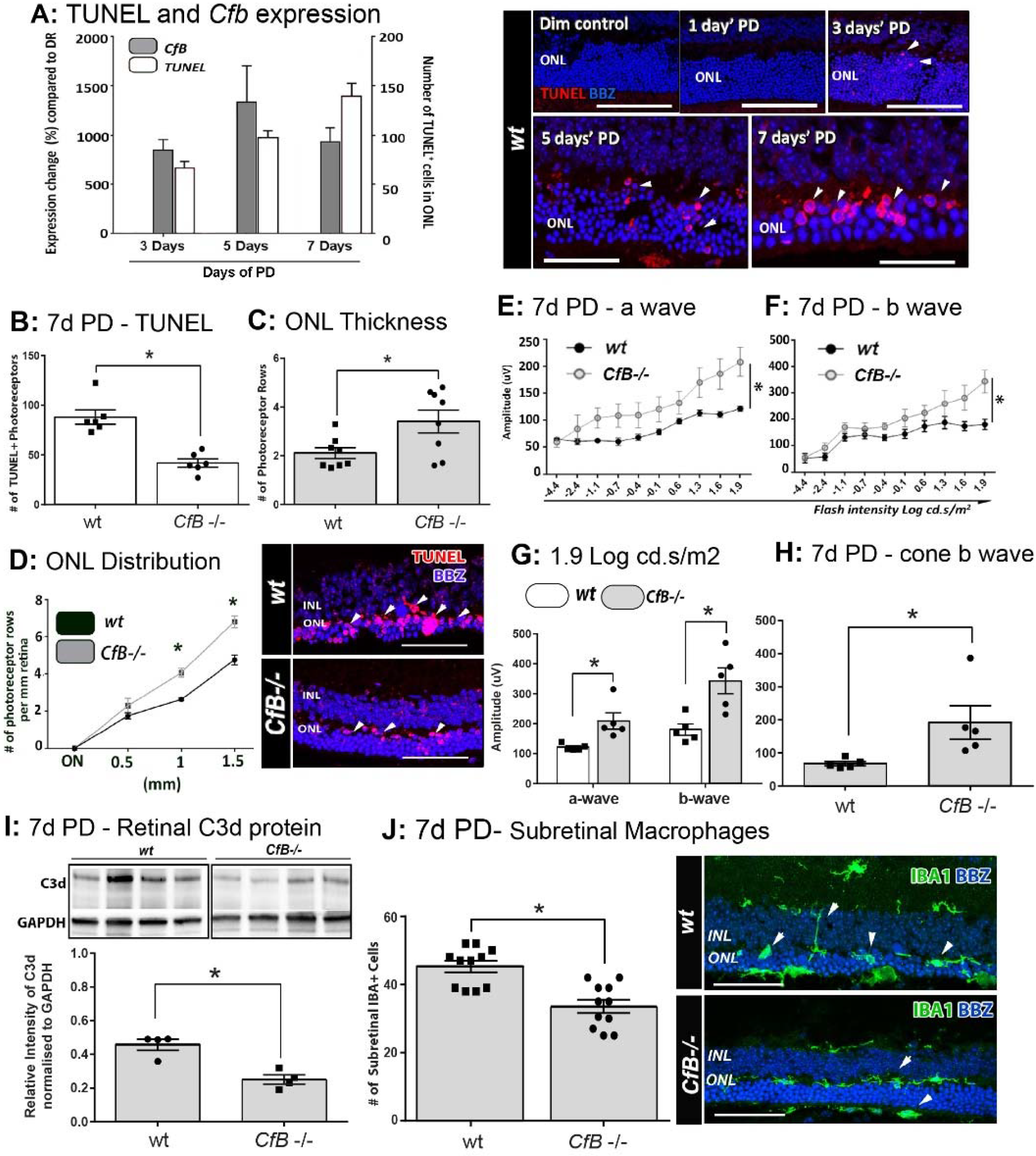
Effect of *Cfb* ablation on retinal degeneration complement activation, and macrophage infiltration following photo-oxidative damage. **A:** Temporal relation of retinal *cfb* expression and TUNEL+ photoreceptor counts PD was assessed in Wt mice after 3, 5, and 7 Days PD. The expression of *Cfb* showed continual up-regulation following PD, compared to dim-reared (P<0.05), and was in concert with increase in TUNEL+ photoreceptors (P<0.05). Representative florescent images showcase TUNEL staining (red) over the timecourse of PD. **B-D:** Change in TUNEL-photoreceptor counts and ONL thickness in Wt vs *Cfb*−/− mice, after 7 days PD. TUNEL+ across the full length of retinal sections were quantified and found to be reduced in *Cfb*−/− mice (B, P<0.05); these are depicted in representative images. ONL thickness was extrapolated on the same sections as function of the number of photoreceptor rows (C-D). When quantified over sections (C), there more surviving photoreceptors in *Cfb*−/− mice, compared Wt (P<0.05), and was particularly pronounced in mid-periphery (~1 to 1.5mm eccentricity, D). **E-H:** ERGs recordings capture a flash intensity series (−4.4 − 1.9 cd.s/ms^2^) conducted on Wt and *Cfb*−/− cohorts after 7 days PD. The trend for both analysed b- and b- waves across this series was higher Cfb mice than Wt (E-F), in addition to highest flash intensity (G, P<0.05). The cone-derived b wave was analysed at from a twin-flash stimulus at 1.9 cd.s/ ms^2^, and was also significantly higher in the *Cfb*−/− cohort (H, P<0.05). **I:** Representative immunoblots illustrate bands for complement C3d protein and loading control GAPDH in whole retinas, from CfB−/− and WT cohorts after 7 days PD. Densitometry quantified C3d levels, normalised to GAPDH, and indicated reduced C3d levels in the *Cfb*−/− cohort (P<0.05). **J:** Immunohistochemistry for IBA1+ macrophages/microglia (green) in retinal sections of Wt vs Cfb−/− mice after 7 days PD, as shown in representative images. The graph illustrates the quantification of IBA1+ cells in the ONL and subretinal space across retinal sections, and show a significant decrease in the Cfb−/− cohort, compared to Wt (P<0.05). Statistical significance was determined by student t-test or ANOVA accompanied with post-hoc multiple comparison (*P< 0.05). A-H: N=5 per group; I-J: N= 4 and 11 per group. ERG, electroretinogram; INL: inner nuclear layer; ONL: outer nuclear layer; PD, Photo-oxidative damage. Scale bars equal to 50μm.

To investigate a possible functional role, we utilised a *Cfb*−/− knockout strain and a murine photo-oxidative damage model that we had previously shown exhibits a similar magnitude of photoreceptor death and sterile inflammation to the rat model, over a 7-day time course of bright-light exposure [43]. We examined the expression of Factor B in whole mouse retinas after 3, 5, or 7 days’ PD, and observed its persistent up-regulation compared to dim-reared retinas (**Figure 7A**; P<0.05). This correlated with increased number of TUNEL+ photoreceptors, following PD over the same period (**Figure 7A**; P<0.05).

The link between photoreceptor viability and complement factor B was further examined in Wt and *Cfb*−/− mice using TUNEL, ONL thickness, and ERG recordings (**Figure 7B-H**). After 7 days’ exposure to PD, fewer TUNEL+ photoreceptors and more surviving photoreceptor rows were observed in *Cfb*−/− mice, compared to Wt (**Figure 7B-D**; P<0.05). In conjunction, ERG analysis indicated significantly better retinal function in the *Cfb*−/− cohort, which had higher mixed a- and b- wave flash responses than the Wt group (**Figure 7E-H**; P<0.05). The status of complement within retinas of *CfB*-/-and Wt was inferred by Western blotting for C3d (**Figure 7I**), a relatively long-lived by-product of C3 proteolysis whose accumulation infers activation of the cascade [58]. After 7 days’ PD, there were significantly lower levels of C3d in retinas of *CfB*−/− mice, compared to Wt (P<0.05), accompanied by fewer infiltrating IBA1-immunoreactive macrophages in the ONL/subretinal space (P< 0.05, **Figure 7J**). IBA1+ macrophages in the Wt cohort more frequently exhibited a reactive amoeboid-type morphology (**Figure 7J**, representative images), indicative of an activated state.

## Discussion

In this study we provide the first detailed characterisation of the functional dynamic of the retinal leukocyte population in sterile retinal inflammation, combining flow-cytometry analysis with RNA-seq of retinal CD45^+^ transcriptome. Whole-genome transcriptional profiling has been used previously to facilitate high-resolution analysis of the host gene response to cellular changes in the retina [59–61], though these studies have lacked the necessary resolution to dissect the leukocyte populations driving tissue damage after sterile injury. Here, we show that the major leukocyte cell type, and associated molecular changes are consistent with a mononuclear phagocyte subpopulation, most likely tissue resident microglia and macrophages. Photo-oxidative damage induced an early pro-inflammatory and chemokine-driven response that laid the foundation for progressive and varied migration of myeloid cells, granulocytes, and T lymphocytes at later stages, which was coincident with photoreceptor cell loss. In assessing the gene expression changes using a functional network approach, we uncovered shifts in the retinal leukocyte transcriptome following sterile injury, predicting the major drivers of the degenerative process are mediated by a sustained inflammatory response. Specifically, early myeloid-driven, acute pro-inflammatory responses preceded any later involvement of complement, T-cell activation, antigen presentation.

Network analysis predicted a role for alternative complement pathway during the late stage of degeneration, and indicated that this was driven by leukocytes at 3-7 days post injury. Factor B is crucial to the assembly of the alternative pathway C3-convertase, which promotes the rapid and prodigious accumulation of C3b/C3d, and ultimately, the assembly of the membrane attack complex (MAC), that in turn may trigger cytolysis or apoptosis of target cells [62, 63]. Factor B expression is increased in retinal macrophages/microglia in aged mice [64], though the implications of this expression, particularly in the context of sterile inflammation, has not previously been assessed. Here, we present several lines of evidence indicating that increased expression of Factor B promotes local activation of the alternative complement pathway, including deposition of complement C3, and that this is mediated by subretinal macrophage infiltration.

AMD is considered a chronic inflammatory disease exacerbated by dysfunction and dysregulation of the complement cascade (reviewed in [65, 66]). Much attention has been placed on the regulation of Complement Factor H (*Cfh*), and thus the alternative pathway (reviewed in [67]), due to the strong genetic link between Cfh dysregulation and AMD pathogenesis [68–71]. The alternative pathway has been implicated in complement activation and retinal pathology in several animal models (reviewed in [72]). This relates favourably to our previous observations, showing that C3 is expressed by infiltrating mononuclear phagocytes following PD, and that this expression is integral to the pathogenic activation of complement within the retina [73].

It is well known that the major leukocytes in the central nervous system are microglia, the resident macrophages (reviewed in [74]). Previous studies have used CD11b^+^ as a leukocyte marker for isolating retinal macrophage populations [75], and have shown that this population accounts for 5-20% of the glial population [76]. Despite CNS microglia being classed as CD45^lo^ expressed [77, 78], in the current study we found that CD45 labelling produced a similar profile to CD11b, but comprise a broader pool of cells, therefore providing a better perspective on the leukocyte expressome. Our data shows that while in control animals (reared in dim light) CD11b^+^ cells are ~0.495% (^+/−^0.09 SEM) of the retinal population, CD45^+^ cells comprise ~0.853% (^+/−^0.23 SEM). However, at 7 days after light damage CD11b+ cells were 5.07% (^+/−^0.530 SEM) while CD45+ cells 6.21% (^+/−^0.730 SEM) of the retinal cell population, respectively. The data indicates that the molecular profile of the CD45^+^ cell population changes with retinal degeneration. Moreover, the early expression of chemokines suggests that the increase in the proportion of CD45+ cells is due to a recruitment of monocytes as well as a change in the local retinal environment.

Previous studies have demonstrated that recruited macrophages do not contribute to the microglial pool under normal physiological conditions [79]. However, recent evidence from Ma and colleagues in an NaIO3-induced model of RPE loss indicates that recruited macrophages may aid in replenishing the microglial pool of the inner retinal during injury. In that study, recruited macrophages were found to seed the inner retina in a Ccr2-dependant fashion following RPE degeneration, and adopt a long-lived phenotype synonymous with resident microglial cells [80]. Our gene expression data support this finding in showing that microglia-associated markers transmembrane protein 119 (*Tmem119*), P2Y purinoceptor 12 (*P2ry12*), and Sal-like protein 1 (*Sall1*) are all transiently down-regulated in the CD45+ population, at 0 days (**Supplementary Table S1**), suggestive of a dilution of the microglial pool with the rapid recruitment of non-resident mononuclear phagocytes. Despite these collective findings, the dissection of the respective functional roles of resident vs recruited macrophage populations in the retina remains elusive. Fate mapping strategies, such as that reported recently by O’Koren and colleagues [78] may further assist in contrasting the roles these populations in future studies.

Network analysis of leukocyte gene expression also highlighted waves of chemokine mediated cell recruitment. Chemokines play a pivotal role in leukocyte migration and activation [81] and are implicated in experimental models of retinal degeneration [47, 82] and in AMD disease progression [29, 83]. These small molecules are grouped according to the relative position of their first N terminal cysteine residues, into C (γ chemokines), CC (β chemokines), CXC (α chemokines), and CX3C (δ chemokines) [23, 84, 85]. Many of the receptors show a degree of redundancy, although generally interactions are restricted to within chemokine family subclasses [23]. We find significant changes in expression of a number of chemokines by CD45+ cells in retinal degeneration, covering the gamut of myeloid and T lymphocyte chemotaxis. The significant increases in *Ccl2*, *Ccl3*, and *Ccl7* at 0 days corroborate findings from our previous studies of whole retinas [30, 47]. Here we show in addition that *Ccl12*, *Ccl17*, *Ccl22*, as well as *Cxcl4*, *Cxcl10*, *Cxclll*, *Cxcl13*, *Cxcl16* are expressed by leucocyte subsets in the retina at early stage of degeneration. The chemokine expression profile at 3-7 days also indicated some striking novel patterns, including strong upregulation of *Cxcl9*, *Xcl1*, *and Xcr1* axis. In a previous study, a downregulation of *Cxcl9* was observed in IFN-B-treated RPE cells, and suggested to be an immune-suppressive mechanism that protects the retina from excessive inflammation [86]. The ligand *Xcl1* is expressed primarily by activated CD8+ T cells in peripheral blood, while its cognate receptor *Xcr1* is present mainly on dendritic cells (reviewed in [87]). This signalling axis augments T cell survival and promotes cytotoxic immune responses [88, 89].

Our network analysis showed that CD45+ expression profile at the later stage of sterile inflammation heavily skewed towards antigen-presentation and processing via MHCI and MHCII, as well as activation and signalling through the T cell receptor. In human forms of sterile retinal inflammation, such as AMD, the evidence linking disease progression to the adaptive immune system has been poorly investigated, even though the presence of anti-retinal antibodies in AMD patients has been reported [90–92]. Others have suggested that AMD should be considered an autoimmune disease (reviewed in [93]), noting that evaluations of AMD lesions demonstrate the presence of both mast cells and lymphocytes [94]. Because the retina is not routinely surveyed by B-and T-cells under physiological conditions, any involvement of the adaptive immune system to retinal degenerations will most likely involve retinal antigen presentation and indirect autoantibody function [42]. T-cells and neutrophil participation in RPE degenerations has also been reported [95, 96], while others have implicated the complement system as the bridge between the adaptive and innate immune system [97, 98], leading to the recruitment of γδ T-cells [99]. A study that induced AMD-like retinal degeneration through the use of carboxyethylpyrrole-modified albumin (CEP) [100], suggests that an antibody-mediated response to CEP is required to initiate degeneration, and implicates T-cells and B-cells. Further, this CEP-immunised AMD-like model demonstrates macrophage recruitment to the site of injury and complement activation in the Bruch’s membrane, also suggesting activation of the classical pathway [100]. The study also shows that CEP-immunised Rag ^−/−^ mice, which lack intact adaptive immunity and mature T-cells and B-cells, produced no anti-CEP.

## Conclusion

Sterile inflammation punctuates the degenerative process of many ocular pathologies, though despite this the breadth and scope of this response in the context of the retinal environment are poorly characterised. Though mononuclear phagocytes comprise the bulk leukocyte infiltrate, profiling the CD45+ subset did reveal an early and pronounced enrichment of terms pertaining to T-cell chemotaxis and migration, while proliferation, T-cell activation, antigen presentation, and complement dominated ontologies at the later time points. Finally, our mechanistic data strongly support a key role of leukocytes, and in particular mononuclear phagocytes, in propagating the subretinal inflammation and complement deposition via the activation of the alternative pathway. Together, these data greatly extend our understanding of the factors that shape the course of sterile retinal inflammation, which has relevance to the therapeutic targeting of these pathways in diseases such AMD.

## Acknowledgments and funding

We would like to thank the members of the ANU Bioinformatics Consultancy Unit, JCSMR, for their help in the analysis of the RNAseq data as well as the funding agency Retina Australia for supporting this work.

